# Structural variation in hippocampal dentations among healthy young adults

**DOI:** 10.1101/2020.02.09.940726

**Authors:** Julia ten Hove, Jordan Poppenk

## Abstract

Little is known about hippocampus dentations – or “bumps” on the inferior surface of the hippocampus – and whether they hold functional significance. Empirical work on these features is beginning to emerge, but no study has yet been conducted in which their spatial positions are demarcated, and their putative role in individual differences in memory is based on a single small study. We gathered ultra-high resolution isotropic MTL scans from young-adult sample and performed such measurements. On the long axis, we found smaller and denser dentations moving towards the hippocampus’ posterior extent. Although other reports have distinguished the functional characteristics of anterior and posterior hippocampal dentations, we found hardly any were actually localized in the hippocampal head, with half of participants featuring no anterior dentations at all. We also observed more numerous dentations in the left hemisphere than right, and found this asymmetry tracked right-hand dominance. All dentations lay specifically within in the CA1 subfield, which was thicker at dentations – protruding below standard lower boundary of the hippocampus – as well as larger in participants with more dentations, suggesting dentations reflect CA1 growth rather than mere folding. Finally, we observed an association between dentations and visual source memory, but not recognition memory. The current report is the first to characterize the spatial properties of this intriguing neuroanatomical feature, which requires the direct spatial labeling of dentations. Further to better characterizing this intriguing neuroanatomical feature, we provide some confirmatory evidence for a link to episodic memory, although we found evidence for this link to be less overwhelming than initially reported.

## 1. Introduction

The hippocampal formation has been the subject of more studies than any other subcortical brain structure, and many brain disorders have been correlated with abnormalities in its volume, shape and metabolic properties (Chang et al., 2018). In spite of this exhaustive work, much less is known about one of its more peculiar anatomical characteristics: the series of bumps sometimes observed along the inferior and lateral surface of the hippocampus. These are believed to arise specifically from the folds of the CA1 subfield and subicular region (Chang et al., 2018). Simic and colleagues (1997) were the first to qualitatively describe the bumps or ridges as “gyrations” of these subfields, and neuroanatomical studies have come to refer to them as dentations.

Early work on dentations has focused on methods for evaluating them, including means of measuring dentations in lower resolution clinical scans (Chang et al., 2018) or conducting software-based characterization of a single individual (DeKraker, Lau, Ferko, Khan, & Kohler, 2019). Qualitative work of this kind has offered a helpful overall picture of dentations, yet the typical distribution of basic descriptive features in the population, such as their size and positioning, have, remarkably, gone entirely undocumented so far in the hippocampal literature. In fact, to our knowledge, there are only two group studies that have reported any data relevant to hippocampal dentations: one involved trained raters conducting holistic assessments in an *n =* 22 sample spanning a large age range of 20 to 57 years (Beattie et al., 2017). Another employed a novel PCA-based algorithm for feature extraction to estimate dentations from the mid- 20’s to 80’s (Cai, Yu, & Zhang, 2019). Although these studies contribute valuable initial insights, in no case have researchers yet presented or validated results by carefully inspecting and labeling hippocampal dentations. Also, the study by Beattie and colleagues drew on a very small sample, whereas the Cai and colleagues’ study relied on low-resolution data. Both datasets also focused exclusively on frequency and amplitude and omit coordinate data or direct measurements of dentation height and width. Finally, neither investigation established relevant values for a healthy, normally developed cohort that is not yet affected by the possible effects of aging-related atrophy.

For further progress in this area, a baseline assessment of healthy young adults is needed, using high-quality brain images and manual inspection. Such a study should incorporate direct measurements, rather than rater summaries or algorithmic estimates. To conduct such an investigation, we gathered high-quality 0.5 x 0.5 x 0.5 mm resolution brain-images, the highest resolution used in a group study to date. More typically, high resolution hippocampal sequences feature high in-plane (typically coronal) voxel dimensions at the cost of thicker slices, in which dentations, which proceed in the same sagittal direction as the thick slices, are obscured. We also gathered images with T2-weighting, in which dentations are more visible than a standard T1-weighting (Winterburn et al., 2015). We sampled these images from a healthy cohort of 66 people, drawing on a narrow age range of 22 to 35 years to avoid developmental effects related to either the final stages of brain maturation or the early stages of brain atrophy (after Van Essen et al., 2012).

3D segmentations of the hippocampus typically show the inferior surface to be smooth or mildly irregular as a result of smoothing or group-averaging in the surface rendering pipeline, factors that are prone to overwhelming the prominence of 3D dentations (Chang et al., 2018), but the high resolution of our input data allowed us to minimize such smoothing effects to identify and label dentations on 3D hippocampal models. By localizing dentation peaks and their neighbouring “valleys” in space, it is possible to directly measure their basic features of height, width, and position, and to relate their position to other features of the hippocampus. In the current case, the high-resolution images support integration of our dentation analysis with subfield labeling and enable us to not only evaluate the gross position of dentations within the hippocampus, but also to evaluate the relationship between dentations and specific subfields.

Further to basic measurement of the hippocampal dentations, initial efforts to evaluate their functional relevance have been conducted. Initially, scientists postulated dentations to be indicative of cortical atrophy, or some form of brain maldevelopment (Giacomini, 1884). More recently, they have been considered a potentially positive adaptation: just as gyrification in the cerebral cortex is associated with higher IQ scores (Luders et al., 2013), it has been proposed that since hippocampal dentations also reflect a type of gyration (DeKraker et al., 2019), they may, therefore, also reflect a functionally relevant structural adaptation (Chang et al., 2018). This idea is possible to test through exploration of individual differences, since dentations have, qualitatively, been shown to be highly variable across individuals (Cai et al., 2019). In light of the known importance of the hippocampus for episodic memory (Moscovitch, Cabeza, Winocur, & Nadel, 2016) including known contributions of the CA1 subfield and subicular region to episodic memory function (Bartsch, Dohring, Rohr, Jansen, & Deuschl, 2011; Farovik, Dupont, & Eichenbaum, 2010), initial exploration of possible functional relevance for episodic memory has focused on that topic. In particular, (Beattie et al., 2017) identified such links between dentations and various measures of episodic memory to assess this putative functional role. As dentations best predicted recognition memory in their study, we incorporated recognition memory measures into our study, as well, complementing these with source memory measures of episodic memory.

To summarize, we designed an experiment intended to characterize hippocampal dentations using a direct labelling approach to measure dentations. Structural variability was assessed using high quality brain images gathered from an age-restricted group of young adults, to characterize dentation properties within this population. To assess the functional relevance of dentations, we supplemented our anatomical assessment with verbal and visual recognition and source memory measures. Due to recent findings that dentations predict a wide set of individual difference measures in episodic memory, we anticipated dentations would similarly predict these memory measures in our own sample.

## 2. Methods

### 2.1. Overview

Ultra-high-resolution neuroimaging scans and memory data were acquired from a sample of 66 healthy young adults. Two raters applied direct and holistic evaluations of dentation measurement to the first 10 participants to assess the reliability of each approach. Hippocampal dentation values were compared with verbal and visual recognition test scores to explore the relationship between hippocampal dentations and memory in healthy adults. Because the neuroimaging data we inspected were gathered as part of a larger study, some aspects of our neuroimaging methods described here are shared with prior reports (Sunavsky & Poppenk, 2019).

### 2.2. Participants

Participants were recruited in Kingston, Ontario, Canada using posters, Facebook, advertisements and web posts on Reddit, Kijiji and Craigslist. Inclusion criteria required all subjects to be between 22-35 years of age (to avoid potential developmental effects), right-handed, an English native-speaker, and to have normal or corrected-to-normal vision and hearing. Exclusion criteria required all subjects to have no contraindications for MRI scanning, have no history of neurological disorders, or recurrent mental illness requiring medication, and the use of any psychotropic medication. Recruits who reported meeting the specified requirements were invited to attend a two hour in-person screening session. At the screening session, demographic eligibility was confirmed, and a simulated MRI scanner was used to rule out claustrophobia, and the inability to keep still (defined as falling below the 95^th^ percentile of low-frequency motion as measured in a reference sample), and participants who did not meet a low baseline value in cognitive tasks including reading and recognition memory. Cognitive baseline was assessed by participants’ ability to respond to at least 33% of encoding trials in a recognition memory task, to obtain a d-prime of at least 0.1 in a subsequent memory test, and to demonstrate acceptable reading comprehension and speed (a TOWRE score of at least 26.4 and a Nelson-Denny score of 2). Participants were excluded for multiple no-shows, failing to follow instructions, rudeness and regular use of illegal drugs. Participants were told they would be compensated CAD$20 for their time (or a pro-rated amount in the case of early withdrawal).

Of the original sample, 67 participants met all the criteria and 66 returned to complete the full experiment. The average age of participants was 26.6 years (SD=4.30 years). Participants identified themselves as men (n=29), women (n=36), and other (n=1). G*Power 3.1 software was used to calculate a sensitivity power analysis: given our sample of n = 66, we had a power of 0.997 to observe a correlation of *r* = 0.51 (the strength of the relationship between dentations and episodic memory by Beattie et al., 2017) using a confirmatory one-tailed test. We also had an 80% chance of confirming any medium-sized effect of *r* = 0.30 or larger.

### 2.3. MRI acquisition

To help participants learn to remain still during their ultra-high-resolution MRI scans, which are susceptible to motion artifact, the day prior to scanning, participants completed a motion biofeedback session. In this session, they were loaded into the same simulated MRI scanner described above and asked to watch a 47-minute nature documentary with MRI sounds overlaid on its soundtrack. At the beginning of the session, they were reminded of the importance of keeping still in the MRI scanner, and asked to use the session as an opportunity to practice this. As the movie played, their head movements were displayed in the form of a bar graph, and an algorithm caused the film to pause for three seconds when head motion exceeded an adaptive threshold, with annoying visual and audio static played, instead.

The day of MRI scanning, brain images were acquired using a 3 Tesla (3T) whole-body MRI scanner (Magnetom Tim Trio; Siemens Healthcare) located in the Centre for Neuroscience Studies at Queen’s University. Over the course of a 1.5-hour scanning session, a variety of sequences were completed. The sequence of greatest relevance to the current investigation was an ultra-high resolution T2-weighted slab, with its volume centered on the medial temporal lobes (MTL; resolution 0.5 x 0.5 mm^2^; 384 x 384 matrix; slice thickness 0.5 mm; 104 transverse slices acquired parallel to the hippocampus long axis; anterior-to-posterior encoding; 2 x acceleration factors; TR 3200ms; TE 351ms; variable flip angle; echo spacing 5.12ms). This protocol was modelled after that used by Chadwick and colleagues (Chadwick, Bonnici, & Maguire, 2014). In addition, whole-brain T1-weighted and T2-weighted whole brain images were acquired and registered to both the MTL slab and a spatial template for the purpose of indexing slab coordinates in standard space (in-plane resolution 0.7 x 0.7 mm^2^; 320 x 320 matrix; slice thickness 0.7 mm; 256 AC-PC transverse slices; anterior-to-posterior encoding; 2 x acceleration factor; T1w TR 2400 ms; TE 2.13 ms; flip angle 8°; echo spacing 6.5 ms; T2w TR 3200 ms; TE 567 ms; variable flip angle; echo spacing 3.74 ms). As the data collection effort involved collaboration with many researchers investigating a variety of individual differences within the participant group, the participants also underwent functional MRI scans and further experimental sessions involving objectives unrelated to the current study goals.

### 2.4. Dentation assessment

Only the inferior surface of the hippocampus was analyzed for dentations. The bumps or ridges that are observed on the superior aspect of the hippocampus are considered to be digitations. These correspond to a different anatomical feature that features little variance among healthy adults (Schultz & Engelhardt, 2014). Accordingly, these were not measured or analyzed in the current study, which focused specifically on dentations. Denoised version of the ultra-high resolution MTL slabs were used for this analysis (Manjon, Coupe, Buades, Collins, & Robles, 2010).

#### 2.4.1. Direct dentation labeling

To annotate dentations within our ultra-high resolution structural images, we first applied the HIPS segmentation tool to MTL images (Romero, Coupe, & Manjon, 2017). Segmentations were created for each participant and the subfields were demarcated by different colours. The segmentations were inspected using ITK-SNAP, a software program that produces 3D renderings of segmentations (Fig. 1), for visual guidance.

**Figure 1.**
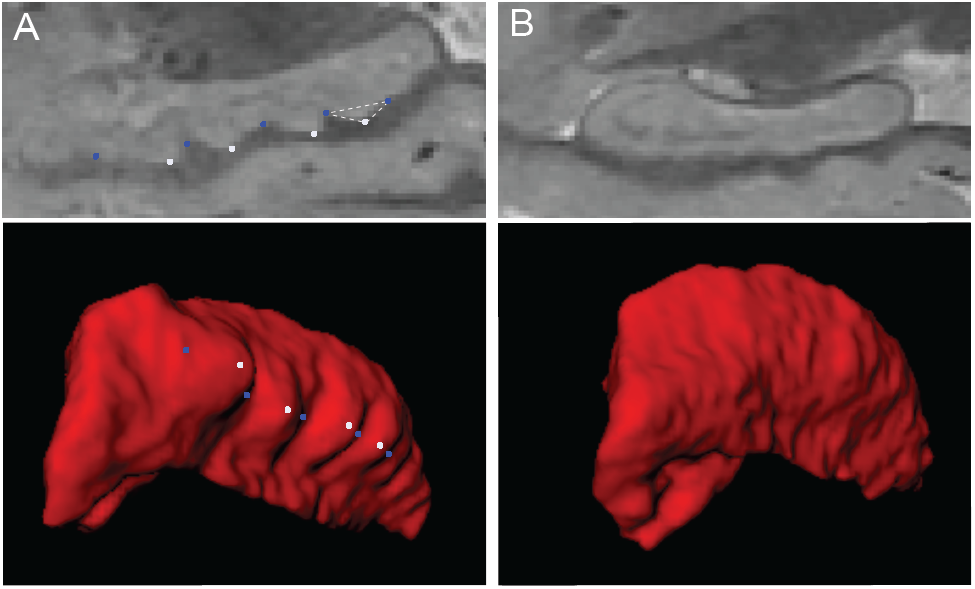
Identification of dentation peaks and valleys using 3D renderings of segmentations. Examples of variation in degree of hippocampal dentations among healthy adults. **A** depicts a hippocampus with numerous dentations on the inferior surface. The top panel is an example source MRI scan, and the bottom panel is a 3D segmentation. Blue labels mark dentation valleys and white labels mark dentation peaks. The dotted triangle represents the geometry used to evaluate dentations using their peaks and two adjacent valleys. **B** depicts a hippocampus that raters deemed to have no dentations. As most dentations did not fall within the same sagittal slice, the overlay on the top row source scans is for illustration purposes.

Using 3D renderings of hippocampal segmentations (smoothed at a default value of 0.8 *SD*), raters identified dentation peaks. For each peak, a corresponding “valley” was also identified, and the last peak on any given hippocampus was associated with an additional (closing) valley such that each peak was bounded by two valleys. The peaks of each dentation were labelled with one value/colour, while valleys were labelled with another. We then algorithmically computed the height and width of the triangle obtained by joining each peak with its nearest two valleys and logged the position of each peak.

Dentations were binned into small and large sizes based on height. Height was calculated first by measuring the length between adjacent valleys to solve for width of the dentation. The width of the dentation with the corresponding peak label was used to solve for area of the dentation. The area and width were then used to solve for dentation height. Our threshold for dentation size was 0.25mm, a value we based on being one half the height of one voxel in the source scan. Based on preliminary inspection of the distribution of dentation heights in our data, we also established a height of 1.5mm as a threshold for a “large” dentation, as this approximately corresponded to the mean height of our sample aligned intuitively with a value equivalent to the height of 3 voxels. Total number of small and large dentations, as well as median height of dentations surviving the minimum threshold, was calculated for each participant.

In follow-up analyses, we evaluated which subfield was closest to each peak by computing its distance to all hippocampal voxels and discovering the subfield label associated with the closest hippocampal voxel. We then evaluated the distance of each peak and valley from the lower bound of the MNI space hippocampus by transforming each peak and valley coordinate to standard space (using a linear transform matrix obtained using Freesurfer) and then, for each point, recording the distance along the z-dimension from the closest point on the standard space hippocampus falling within the same coronal slice. We also computed thickness for each peak and valley coordinate by evaluating the standard space distance, in mm, between each coordinate and the top of the CA1 layer (in the column directly above the coordinate).

#### 2.4.2. Holistic scoring of dentations

The holistic evaluation method of Beattie and colleagues (2017) was adopted. The rating scale incorporates both quantity and prominence of dentations in the posterior region of the hippocampus. Regarding quantity, dentations were binned in to four or more dentations (two points), one to three dentations (one point), and no visible dentations (zero points). Prominence of the dentations was characterized as arciform (two points), sinusoidal (one point), or absent (zero points). These dimensions were then integrated into an overall holistic score for each hippocampus, calculated as the square of that hippocampus’ quantity score added to the square of that hippocampus’ prominence score.

### 2.5. Analysis of brain data

To assess IRR for the two forms of rater evaluation (direct measurement and holistic measurement), an intra-class correlation coefficient (ICC) was calculated for each set of dentation measurements using scores from the two raters. All values between both raters were compared for consistency in a two-way random model. Direct labeling performance by raters was evaluated using total quantity of dentations, and was found to be very high, ICC = 0.924. Holistic assessment performance by raters was assessed using overall scores for each participant (taken as a sum across hemispheres, following Beattie et al., 2017) and was also very high, ICC = 0.921.

Next, descriptive statistics were then computed to characterize the quantity and prominence of dentations along the longitudinal axis of the hippocampus and between each hemisphere. Overall dentation quantity was compared to performance scores on visual and verbal recognition tests for each participant to explore the relationship between hippocampal dentations and episodic memory function. Results were computed using Matlab 2018a software.

To evaluate the reliability of each test statistic, bootstrap resampling was employed to establish 95% confidence intervals around the relationship around each variable or between each pair of variables. The use of bootstrap-based statistics provides maximal statistical power when working with smaller samples and avoids making normality assumptions associated with parametric statistics (McIntosh & Misic, 2013). Briefly, 1000 bootstrap samples were obtained, with each representing a random resampling with replacement of the original group, and a correlation coefficient was recorded from that bootstrap sample. Where dependent measures were pooled (e.g. with all dentations being entered into long-axis analysis), resampling was still conducted on a subject-wise basis, with all entries corresponding to a particular participant being entered at once with replacement (i.e., “super-subject” analysis; see also Detre, Natarajan, Gershman, & Norman, 2013). The bootstrap standard error was established by evaluating the variance across samples and a bootstrap ratio (BSR) was computed in which the correlation obtained from the full group was divided by the bootstrap standard error, yielding a value analogous to a z-score that could be used to look up a corresponding *p*-value. Correlations were considered reliable only when this *p*-value fell below 0.05 and their 95% confidence intervals did not encompass zero.

### 2.6. Memory testing

Participants completed a verbal recognition test (dog names) and a visual recognition test (pictures of cats) in separate testing sessions. Other than different materials, the tests shared the same properties. This recognition testing involved an encoding and retrieval phase separated by a distractor task. Both sessions were conducted in an isolated computer testing room booth using a desktop computer running Ubuntu (14.04 LTS). Tasks were executed using Matlab 2014b, Psychtoolbox (Kleiner et al., 2007) and SuperPsychtoolbox (Mountjoy & Poppenk, 2015).

Prior to encoding, participants were advised they would be viewing items under different kind of instructions. For some items, they would be required to guess the dog/cat’s sex (using the M or F keys for male or female, respectively), and for others, whether it seemed like a more sporty or lazy pet (using the S or L keys, respectively). In total, 60 items were presented for each recognition test, and half were presented under each task, with a randomized assignment of task and stimulus sequence. Each item was presented for four seconds, with a 0.75 s inter-stimulus interval. Four additional practice items were studied prior to the main set. After the encoding phase, participants were presented with a distractor task lasting one minute. During this time, they were presented a number, and attempted to type the value of the number minus three. When they pressed enter, the presented number was updated if they were correct, and the task continued, such that they were able to make as many successful responses as possible.

Recognition testing followed the distractor task. All of the original 60 items, plus an additional 60 lures, were presented with no response deadline. Four practice targets and four practice lures were also presented before the main set. For each item, participants first evaluated whether each item was a remember, know, or new item (see e.g., (Dudukovic, 2005). Next, they were asked whether they performed the male-female vs. sporty-lazy task on that same item. Performance was evaluated by 1) collapsing remember and know responses into an “old” response bin and using this to compute d’ for each participant; and 2) evaluating the proportion of correct source responses.

## 3. Results

### 3.1 Demographic factors

Perhaps because they all reflected inter-related characteristics of the same anatomical structures, our dentation measures were mostly correlated with one another (Fig. 2A). The exception was width, which was correlated only with height. However, none of the relationships were deterministic (all *r*’s < 0.81), suggesting they addressed divergent qualities of dentation variance. Holistic ratings were the most correlated with dentation count, suggesting it is numeracy of dentations that typically dominates this metric.

**Figure 2.**
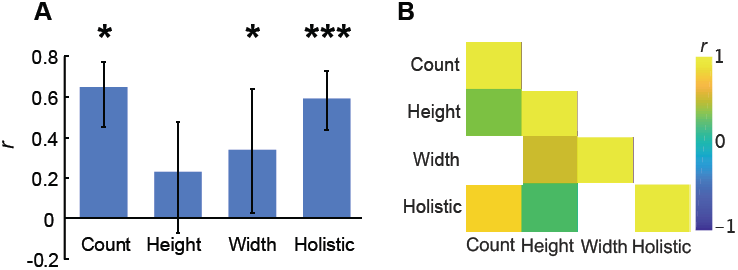
Correlations among dentation measures. A bar graph compares total dentations, dentation width, height, and holistic scores between the left and right hemisphere (**A**). A correlation matrix compares dentation quantity, height and width with holistic scores (**B**). Correlations not significant at *p* < 0.05 in this matrix are suppressed (white).

We attempted to explain this variance as a function of participants’ basic demographics but observed null effects for all analyses. There were no sex differences in overall dentation count between men (*M* = 7.00, *SD* = 4.07) and women (*M* = 8.00, *SD* = 3.94), BSR = −1.02, 95% CI = [-2.89 0.88], *p* = 0.307; in dentation height between men (*M* = 1.25 mm, *SD* = 0.28 mm) and women (*M* = 1.20 mm, *SD* = 0.30 mm), BSR = 0.76, 95% CI = [-0.09 mm 0.21 mm], *p* = 0.448; in dentation width between men (*M* = 5.73 mm, *SD* = 1.69 mm) and women (*M* = 6.09 mm, *SD* = 2.02 mm), BSR = −0.75, 95% CI = [-1.36 mm 0.59 mm], *p* = 0.453, nor in holistic assessments of dentations between men (*M* = 3.53, *SD* = 2.53) and women (*M* = 4.07, *SD* = 2.71), BSR = −0.83, 95% CI = [-1.79 0.34], *p* = 0.406. With regards to age, we found no association with total number of dentations, *r* = 0.06, 95% CI = [-0.18 0.27], BSR = 0.49, *p* = 0.625; with dentation height, *r* = 0.14, 95% CI = [-0.09 0.36], BSR = 1.30, *p* = 0.209; with dentation width, *r* = 0.16, 95% CI = [-0.07 0.36], BSR = 1.40, *p* = 0.152; nor with holistic dentation ratings, *r* = 0.14, 95% CI = [-0.10 0.36], BSR = 1.20, *p* = 0.245.

### 3.2. Gross topography

We began our characterization of dentation locations by examining possible hemispheric effects. Dentation count, height, and holistic scores were correlated between hemispheres, although no relation was found in dentation width between left and right hemispheres (Fig. 2B). However, we did observe a leftward bias, with participants having more dentations overall in the left than right hemisphere, *M* = 0.72, *SD* = 1.88, BSR = 3.07, 95% CI = [0.30 1.21], *p* = 0.002 (Fig. 3). This effect was not apparent in small dentations, BSR = 0.31, 95% CI = [-0.47 0.68] *p* = 0.758, but was apparent in a larger number of large dentations in the left than right hemisphere, BSR = 3.08 95% CI = [0.24 1.06], *p* = 0.002. The same effect was similarly expressed in median dentation height, BSR = 2.30, 95% CI = [0.03 0.31], *p* = 0.021, as well as holistic measurement score, BSR = 2.15, 95% CI = [0.15 2.02], *p* = 0.031. There was, however, no difference in terms of median dentation width between the left hemisphere (*M* = 6.21 mm, *SD* = 2.87 mm), and the right (*M* = 6.17 mm, *SD* = 2.41 mm), BSR = 0.08, 95% CI = [-0.83 mm 0.94 mm], *p* = 0.936. Using the MNI coordinates of dentation peaks along the y-axis, we next explored the position of dentations along the longitudinal axis of the hippocampus (Fig. 4). A trend was found towards more anterior dentations in the right than left hippocampus, BSR = −1.72, 95% CI = [-0.39 0.00], *p* = 0.085, but there were significantly more posterior dentations in the left than right hippocampus, BSR = 2.61, 95% CI = [0.23 1.58], *p* = 0.009.

**Figure 3.**
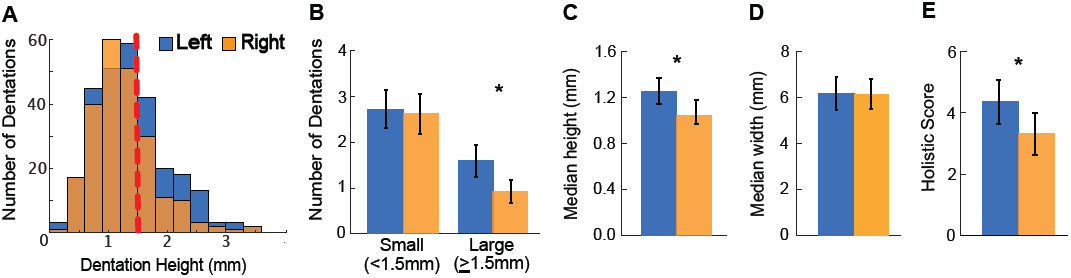
Size and quantity of dentations in left and right hemispheres. A histogram describes the height and number of left and right hemisphere dentations pooled from all participants (**A**). The red dashed line marks the boundary used to distinguish small dentations (<1.5mm) from large dentations (1.5mm) in subsequent panels. **B** reveals an interaction in hemispheric asymmetry as a function of size. **C, D** and **E** compare median dentation height, median dentation width and holistic score ratings for each hemisphere, respectively. Error bars reflect 95% bootstrap confidence intervals. * designates comparisons significant at *p* < 0.05.

**Figure 4.**
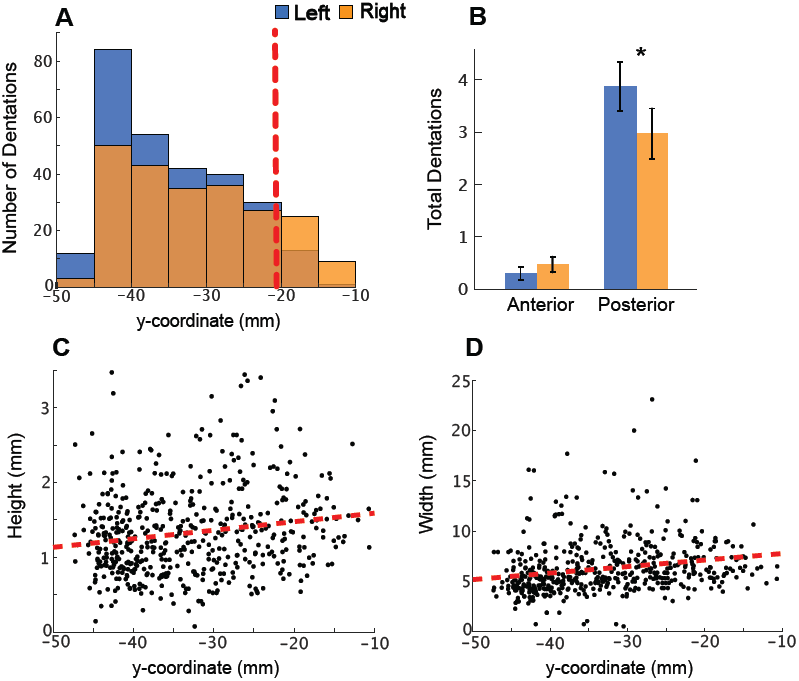
Distribution of dentations along the hippocampal long axis. A histogram describes the distribution of all pooled dentations as a function of position within the y-plane along the hippocampal long axis (**A**). The red dotted line marks the approximate position of the uncal apex landmark used to distinguish anterior from posterior hippocampus. More negative values correspond to more posterior positions along this axis. Dentations in the left hemisphere were located more posteriorly. **B** compares anterior and posterior dentation quantity for each hemisphere. **C** and **D** depict dentation height and width moving from the hippocampus’ posterior to anterior extent (i.e., increasing y-coordinates). Error bars reflect 95% bootstrap confidence intervals. * designates comparisons significant at *p* < 0.05.

Absent any other explanation for these asymmetries, we speculated they could have arisen from the fact that, as in many neuroimaging studies, we had incorporated right-handedness as one of our criteria for inclusion in our experiment, which is associated with left-hemispheric dominance (Knecht et al., 2000). Although we lacked comparative data from left-handed individuals to test this idea, we gathered Edinburgh handedness inventory scores (10-item variant; Oldfield, 1971) as part of our demographics questionnaire, which allowed us to relate the *degree* of right-handedness with number of dentations in a series of post-hoc tests. Consistent with this idea, we found that Edinburgh handedness scores (where larger scores correspond to greater right-hemispheric dominance) were positively correlated with having more right than left dentations, *r =* 0.28, BSR = 3.30, 95% CI = [0.10 0.44], *p* < 0.001; as well as greater holistic scores for the right than left hippocampus, *r =* 0.34, BSR = 3.50, 95% CI = [0.14 0.50], *p* < 0.001. This pattern was not, however, reflected in greater right than left dentation height, *r =* -0.03, BSR = −0.23, 95% CI = [-0.30 0.26], *p* = 0.819, nor greater right than left dentation width, *r =* 0.01, BSR = 0.05, 95% CI = [-0.31 0.27], *p* = 0.957.

Left and right hemisphere data were collapsed to simplify subsequent analysis and to reduce the number of tests performed. A pooled set of all dentations, when moving towards the hippocampus’ anterior extent, grew taller, *r* = 0.17, 95% CI = [0.08 0.26], BSR = 3.71, *p* < 0.001, and wider, *r* = 0.21, 95% CI = [0.13 0.30], BSR = 4.91, *p* < 0.001. In fact, dentations were so sparse in anterior hippocampus that half of participants (32/66) had no dentations at all in the anterior segment for either hemisphere.

To further explore the relationship between dentations and the other aspects of hippocampal anatomy, coordinates of the lower boundary of a standard space hippocampus were identified. For each peak and valley, its z-position was computed relative to the closest point on this hippocampal lower bound falling within the same slice of y. It was found that the peaks fell reliably below the lower bound, *M* = −0.83 mm, BSR = −9.03, *p* < 0.001, but the valleys were not distinguishable in the z-position from the lower bound, *M* = 0.048, BSR = 1.09, *p* = 0.274, and valleys were also significantly higher than peaks, BSR = 8.54, *p* < 0.001 (Fig. 5). A hippocampus upper bound was also computed using a similar approach, and for each peak, compared this upper bound to the top hippocampus voxel from the same slice of y. These “top of the hippocampus” z-positions, too, were indistinguishable from the standard space hippocampus’ upper bound (*M* = −0.043, BSR = −0.55, *p* = 0.580).

**Figure 5.**
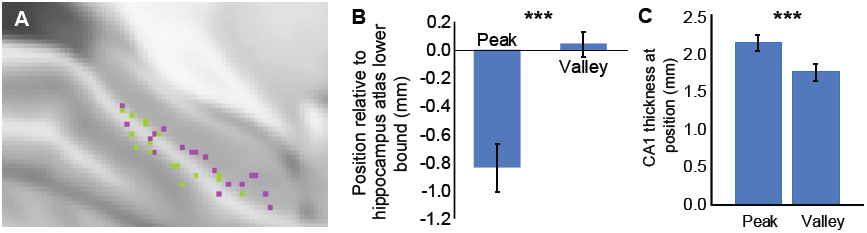
Location of peaks and valleys. Whereas peaks (green dots) typically fell below the lower bound of a standard hippocampus in MNI space, valleys (pink dots) were not distinguishable from this lower bound (**A**), as computed in (**B)**. Thickness of CA1 was significantly greater over peaks than valleys (**C**). Error bars reflect 95% bootstrap confidence intervals. *** designates comparisons significant at *p* < 0.001.

This pattern suggested a specific relationship between dentations and volumetric changes local to the inferior aspect of the hippocampus, consistent with the putative link between dentations and CA1 and subiculum. Accordingly, we sought to determine which hippocampal subfields were closest to each dentation peak. In 100% of cases, dentation peaks fell within the CA1 subfield, appearing to add specificity to prior assessments that dentations fall within CA1 or subiculum. Along these lines, CA1 subfield thickness was greater over peaks, *M* = 2.13 mm, *SD* = 1.26 mm, than valleys, *M* = 1.75 mm, *SD* = 1.24 mm, 95% CI = [0.22 0.53], BSR = 4.80, *p* < 0.001. This evidence suggests that dentations reflect a combination of downward-bound thickening of the CA1 subfield, along with some folding within the CA1 layer.

Based on this proposal, an increase in CA1 volume should be predicted as the number of dentations increases. Consistent with this prediction, we found that a larger number of dentations was associated with a larger proportion of CA1 volume, *r* = 0.38, 95% CI = [0.17 0.57], BSR = 3.77, *p* < 0.001 (Fig. 6). However, no relationship was observed between dentations and overall hippocampal volume, *r* =0.04, 95% CI = [-0.15 0.23], BSR = 0.41, *p* = 0.679, suggesting this change took place at the expense of other hippocampal subregions. Consistent with this idea, more dentations reflect a smaller proportion of CA2/3, *r* =-0.29, 95% CI = [-0.44 -0.14], BSR = −3.73, *p* < 0.001, and a smaller combined stratum radiatum (SR), stratum lucidum (SL) and stratum moleculare (SM) volume, *r* = 0.07, 95% CI = [-0.19 0.33], BSR = 0.49, *p* < 0.001.

**Figure 6.**
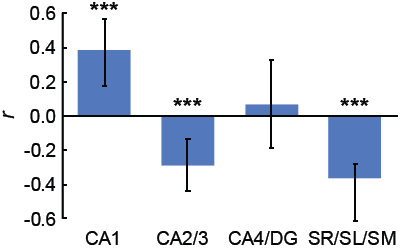
Relationships between number of dentations and subfield volumes. Bar graph comparing total dentation scores correlated to subfield volumes for both the left and right hemisphere. Abbreviations: Cornu Ammonis 1 (CA1), Cornu Ammonis 2/3 (CA2/3), Cornu Ammonis 4 and Dentate Gyrus (CA4/DG), and Stratum Radiatum, Lucidum, and Moleculare (SR/SL/SM). Error bars reflect 95% bootstrap confidence intervals. *** designates comparisons significant at *p* < 0.001.

### 3.4 Relationship between memory and dentations

Beattie and colleagues (2017) have previously illustrated relationships between dentations and episodic memory, with the strongest such relationship expressed between dentations and recognition memory. We attempted to extend this result by attempting to predict both verbal and visual recognition and source memory test scores – both widely recognized metrics of episodic memory - using our dentation measures.

As a first step, we verified that participants’ performance was above chance. We found this to be true for the visual recognition test, as measured by d’ (*M* = 0.60, *SD* = 0.33, chance = 0), BSR = 14.88, *p* < 0.001, and source memory accuracy (*M* = 0.56, *SD* = 0.07, chance = 0.5), BSR = 6.46, *p* < 0.001. Similarly, recognition memory was above chance for the verbal recognition test as measured by d’ (*M* = 1.38, *SD* = 0.38, chance = 0), BSR = 29.71, *p* <0.001, and source memory accuracy (*M* = 0.69, *SD* = 0.09, chance = 0.5), BSR = 17.43, *p* < 0.001. We therefore proceeded to examine how well recognition scores were predicted by dentations.

Broadly speaking, these recognition measures were not associated with number of dentations. There was no relationship between *d’* for visual recognition and total dentations, *r* = 0.00, 95% CI = [-0.25 0.24], BSR = 0.00, *p* = 0.999, dentation height, *r* = −0.05, 95% CI = [-0.31 0.18], BSR = −0.433, *p* = 0.665, dentation width, *r* = 0.05, 95% CI = [-0.11 0.22], BSR = 0.59, *p* = 0.557, nor holistic scores, *r* = −0.08, 95% CI = [-0.33 0.19], BSR = −0.64, *p* = 0.521. Likewise, for verbal recognition, there was no relationship between *d’* for verbal recognition and total dentations, *r* = 0.03, 95% CI = [-0.20 0.27], BSR = 0.23, *p* = 0.816, dentation height, *r* = 0.06, 95% CI = [-0.23 0.30], BSR = 0.43, *p* = 0.667, dentation width, *r* = −0.01, 95% CI = [-0.31 0.25], BSR = −0.06, *p* = 0.951, nor holistic scores, *r* = −0.03, 95% CI = [-0.29 0.21], BSR = −0.28, *p* = 0.780.

Turning to source memory, there was a positive relationship between visual source memory and total dentations, *r* = 0.32, 95% CI = [0.09 0.52], BSR = 2.90, *p* = 0.004, and holistic scores, *r* = 0.24, 95% CI = [0.01 0.45], BSR = 2.18, *p* = 0.030, but only a trend towards dentation height, *r* = 0.28, 95% CI = [-0.03 0.54], BSR = 1.87, *p* = 0.065, and not dentation width, *r* = 0.14, 95% CI = [-0.12 0.38], BSR = 1.15, *p* = 0.251. By contrast, there were no relationships with verbal source memory, including total dentations, *r* = 0.07, 95% CI = [-0.16 0.28], BSR = 0.63, *p* = 0.532, dentation height, *r* = 0.05, 95% CI = [-0.19 0.32], BSR = 0.37, *p* = 0.709, dentation width, *r* = −0.22, 95% CI = [-0.42 0.02], BSR = −1.93, *p* = 0.053, and holistic scores, *r* = 0.00, 95% CI = [-0.27 0.24], BSR = 0.02, *p* = 0.980.

## 4. Discussion

Here we reported a normative analysis of hippocampal dentations among a healthy cohort of young adults. We observed more dentations in the left hippocampi than the right and linked this difference to subtle variations in handedness. We also observed smaller and denser dentations moving towards the posterior extent of the hippocampus along its longitudinal axis, with dentations appearing so sparsely in the hippocampal head that nearly half of participants featured no dentations in either hemisphere. Age and sex did not interact with any property of dentations. We observed dentations exclusively within the CA1 subfield and found the number of dentations to be positively correlated with proportion of CA1. Finally, we observed a link between number of dentations and visual source memory, but did not find link between verbal or visual recognition memory and dentations, as were observed in the only other report assessing dentations as an individual differences measure.

### 4.1. Gross topography of dentations

Dentations were taller and more numerous in the left hemisphere. Beattie and colleagues (2017) observed a numerically larger number of dentations in the left hippocampus, although this difference was non-significant. In the current study – which featured three times the sample size and a more precise direct labeling approach, this pattern was substantiated by nearly all of our variables, with more numerous and larger dentations in the left hippocampus, suggesting the null effect observed in the prior study was a type 2 error arising from low power. In a *post-hoc* effort to explain this lateralization, we speculated that the effect might arise from the fact that we specifically recruited right-handed participants for the current research (corresponding to left hemispheric dominance; Knecht et al., 2000). Consistent with this notion, although our sample was limited to right-handed participants, we found the asymmetry to be predicted by *degree* of right-handedness observed using the Edinburgh Handedness Inventory, with greater right-handedness yielding greater right-leaning asymmetry of dentations. Although this result should be confirmed in subsequent analyses using a cohort that includes left-handed individuals, it reveals a possible link between dentations and hemispheric dominance; however, the nature of this relationship is unclear.

Turning to an anterior/posterior, rather than the left/right dimension, we found dentations to be more numerous in the posterior segment. Using the landmark established in the EADC-ADNI Harmonized Hippocampal Protocol for manually distinguishing anterior from posterior segments of the hippocampus (the uncal apex; Apostolova et al., 2015), we in fact found no anterior dentations at all in the hippocampal head of about half of participants. This raises the question of how Beattie and colleagues (2017) were able to report on different memory effects in their count of anterior and posterior hippocampus dentations. In their study, the anterior/posterior hippocampus was defined by a coronal plane through the quadrigeminal (tectal) plate in the dorsal midbrain, identified for each participant. However, this landmark is located so far posteriorly that it has traditionally been used to distinguish the hippocampus body from the tail (Gan, 2015). Using the more traditional landmark of the uncal apex, dentation count within the head is so frequently zero that analysis of dentations in the anterior may be problematic.

Having made this observation, a natural follow-up question to pose is “where are the missing dentations” in the hippocampal head? It is tempting to turn to the characteristic ridges found on the superior aspect of the head, referred to as digitations. Hippocampal digitations, unlike dentations, are located on the superior aspect of the hippocampal head rather than on the inferior aspect of the body and tail regions, where dentations are readily observed. However, digitations are present among all adults, lacking the normal variation of dentations among healthy individuals (Ding & Van Hoesen, 2015). In addition, they do not appear to consistently arise from the CA1 subfield. An alternate explanation is that the complex organization and folding pattern of the hippocampal head causes CA1-based anterior dentations to be obscured from view. Indeed, much of CA1 in the head is contained in inward folds of the structure. Analysis of histology-scale group data using novel unfolding algorithms (e.g., DeKraker et al., 2019) may be the most promising approach to solving this problem, once technical advances allow for high-resolution neuroimaging data of this kind to be obtained economically.

### 4.2. Relationship of dentations to CA1

To date, dentations have been described as falling in both the CA1 and subicular subfields (Beattie et al., 2017; Chang et al., 2018). A novel contribution of our analysis, therefore, was to use our high-resolution hippocampal images to delineate subfields and evaluate their positions relative to these (and other) subfields. However, whereas we expected to find dentations in both subfields, we found them to be located exclusively in CA1. We also found number of dentations to be positively correlated with the proportion of the hippocampus made up of CA1 and did not see any such relationship between dentation count and subicular tissue. Although this result appears to refine what is known about where dentations typically appear, it is notable that the boundary between CA1 and the subiculum, which neighbour one another on most of the inferior hippocampal surface, may be differently delineated by different protocols. It may, therefore, be possible to obtain a different result using definitions different from those implemented by Winterburn and colleagues (J. L. Winterburn et al., 2013), whose atlases were used to train the HIPS software (Romero et al., 2017) used for segmentation in the current study. Nonetheless, the current results do reveal that, at least according to one prevailing system of subfield definitions, the adherence of dentations to CA1 is more selective than previously thought.

We next attempted to address where dentations fell relative to the typical boundary of the hippocampus. Do dentations reflect protrusions from the hippocampus, or a folding inward? To address this question, we evaluated the position of dentation peaks relative to the normal inferior surface of the hippocampus in standard space. Whereas valleys aligned with the inferior surface, dentation peaks fell reliably below it. Along these lines, we found CA1 to be thicker over dentation peaks than valleys. Combined with evidence that dentations contribute to the relative makeup of CA1 volume, these results suggest that dentations are a feature reflecting a “sprouting” or protrusion from a typical hippocampus; extending inferiorly from its inferior surface towards the para-hippocampal gyrus. It should be noted, however, that this increase in CA1 volume appears to trade off against larger CA2/3 and SL/SR/SM volumes, such that no overall increase in the size of the hippocampus results from the changes.

### 4.3. Dentations and episodic memory

Beattie and colleagues (2017) have been the only authors to date to compare dentation scores in healthy adults to episodic memory function. Based on research that showed a positive correlation between cortical gyri (i.e. increased cortical surface area), and cognitive functioning (Luders et al., 2013), they hypothesized that increased hippocampal dentations will create a larger surface area and therefore correlate to an increased capacity for episodic memory in healthy adults. The best predictor in their study was a measure of visual recognition memory. We therefore tested verbal and visual recognition memory measures and related these to our dentation measures, postulating that hemispheric interactions between test scores and dentations might arise on the basis of stimulus modality. However, we found no relationship between recognition memory scores and any measure of dentations. We obtained this result even when using the same dependent measure (holistic dentation ratings) used by the original authors, and in spite of adequate power arising from a sample size three times that used to identify the original result. In particular, we had power of 0.997 to observe a correlation of *r* = 0.51 using a confirmatory one-tailed test, as well as power of 0.8 for confirming any effect of *r* = 0.30 or larger. Accordingly, it is unlikely that, by chance, we failed to observe a true relationship of the size originally reported that pertains to episodic, or even recognition memory. Rather, because effect-size estimates are more variable around their true value in small samples, it is more likely that the originally reported link between episodic memory and dentations in that analysis exceeded its true value in the study by Beattie and colleagues (*n* = 22).

Although we were unable to detect any relationship between dentations and recognition memory, we did observe correlations between features of dentations (total count and holistic scores) and a source memory measure embedded in the visual recognition memory test, and dentation count. Source memory is widely recognized as drawing upon episodic memory, and has previously been successfully linked to structural properties of the hippocampus (Johnson, 2005; Poppenk & Moscovitch, 2011). However, there is no clear reason to suppose that visual, but not verbal source memory should feature a relationship with dentations. For instance, Beattie and colleagues (2017) observed relationships with memory tests of both verbal and visual modalities. Also, although it is possible that recognition memory in our paradigm relied on non-episodic forms of information (namely familiarity; Yonelinas, 2002) the rate of recollection responses was also unrelated to all dentation measures in both paradigms (all *p’*s > 0.1).

Taking these considerations together, there was no strong evidence to substantiate nor refute the notion of links between hippocampal dentations and episodic memory. Due to the selective nature of our positive visual source memory result in spite of adequate power, we suggest that claims regarding a link between dentations and episodic memory should be interpreted with caution until further work can be conducted. Given the link between dentations and CA1, the functional roles that have been proposed for CA1 may provide a reasonable alternative direction for predicting which memory tasks and measures might be more successfully associated with dentations. Exploring the association between dentations and memory tasks that specifically target this subfield may be a valuable direction to pursue.

### 4.4. Relation between direct labeling and holistic assessment of dentations

In our investigation, both direct labeling and holistic rating procedures yielded high inter-rater agreement. Dentation count and holistic scores were also highly correlated, and perhaps unsurprisingly, we found that all of the measures behaved similarly in descriptive analyses, suggesting they index similar information. We therefore suggest the holistic evaluation may be suitable for evaluation of dentation prevalence where expedience is important, such as in clinical applications; or where a large number of participants are to be evaluated, such as in large datasets. However, where precise measurements or greater breadth of information are required, direct labeling is advantageous for providing an exact dentation count along with height, width, and position information.

### 4.5. Conclusions

In our study, we characterized the spatial positions of hippocampal dentations in a healthy population. Among our observations were observation of smaller but more numerous dentations in the posterior hippocampus, with zero to few dentations in the hippocampal head proper; and more numerous left than right dentations, a pattern we linked to handedness. We also observed evidence linking dentations specifically to the CA1 subfield, as well as evidence suggesting that dentations reflect downward protrusions of that subfield towards the parahippocampal gyrus. Finally, we observed evidence linking dentations to visual source memory, but not other expected forms of memory, suggesting the link between dentations and memory is not is as clear as was once envisioned. Overall, we believe that the current basic characterization of these anatomical features will contribute to a better understanding of hippocampal dentations, and in so doing, contribute to a foundation for research into their role in individual differences.

## Acknowledgements

We gratefully acknowledge Nelly Matorina, Julie Tseng, Natalie Doan, Lauren DeMone, Nigel Barnim, Sarah Berger, Megan Fleming, Maddie Gillis, Sophie Kinley, Lindsay Lo, Lydia Luchich, Jane Mao, and Gillian Marvel for assistance with behavioural data collection. We also thank Julie Tseng, Lauren DeMone, and Natalie Doan with scheduling; Julie Tseng and Don Brien with MRI data acquisition; Justin Siu, Roland Dupras and Mike Lewis with technical support; and Aleksander Biorac for assistance with methodological development and validation. This research was funded by Natural Sciences & Engineering Research Council Discovery Grant 03637 (J.P.). Infrastructure funding was provided by Canada Foundation for Innovation – John R. Evans Leaders Fund (J.P.), and a Queen’s University Research Initiation Grant to J.P., who was supported by the Canada Research Chairs program.

## Data statement

Our analyses were conducted using publicly available software, as described in the manuscript, used together with custom scripts. Neuroimaging scans, cognitive data, and scripts are available upon reasonable request.

